# Ibrutinib modulates CD19 expression and improves efficacy of CD19-CAR T cells in B-cell lymphoma models

**DOI:** 10.64898/2026.05.22.727139

**Authors:** Sisi Wang, Alberto J Arribas, Chiara Tarantelli, Elisa Civanelli, Guangyu Zhou, Pragallabh Purwar, Amandine Pradier, Astrid Melotti, Francesca Guidetti, Yara Hajj Younes, Lodovico Terzi Di Bergamo, Luciano Cascione, Sara Napoli, Emanuele Zucca, Elena Sotillo, Crystal Mackall, Manel Esteller, Yves Chalandon, Davide Rossi, Federico Simonetta, Francesco Bertoni

## Abstract

Antigen density critically influences CAR T-cell efficacy. We show that ibrutinib induces CD19 upregulation in B-cell lymphoma models, enhancing CAR T-cell cytotoxicity. Pharmacologic modulation of target antigen expression represents a promising strategy to overcome resistance.

---

Chimeric antigen receptor (CAR) T-cell therapy has redefined the management of relapsed or refractory B-cell malignancies, yet a substantial proportion of patients with B-cell non-Hodgkin lymphoma (B-NHL) fail to achieve durable responses. While variability in CAR T-cell function, expansion and persistence contributes to heterogeneous outcomes, tumor-intrinsic resistance mechanisms are increasingly recognized as key determinants of treatment failure. Among these, modulation of target antigen expression has emerged as a central mechanism of resistance.

CD19, a lineage-restricted glycoprotein that enhances B-cell receptor signaling, is the protein target of currently approved CAR T cells for B-NHL. Antigen density significantly influences CAR T-cell activation thresholds and cytolytic potency ^1,2^. Surface CD19 levels are often lower in lymphoma cells than those in normal B cells or acute lymphoblastic leukemia cells ^2^, although true CD19-null lymphomas are rare^3^. Loss or downregulation of CD19 after CAR T-cell therapy frequently occurs, with up to 60% of relapsed diffuse large B-cell lymphoma (DLBCL) patients showing this change ^4,5^. However, flow cytometry and molecular analyses indicate that genomic loss is rare ^6,7^, and most cases involve CD19 downregulation rather than total loss. These observations suggest that pharmacologic modulation of CD19 expression could restore CAR T-cell recognition and overcome antigen-dependent resistance. Since Bruton’s tyrosine kinase (BTK) signaling influences B-cell receptor-driven transcriptional programs, we investigated whether BTK inhibition could increase CD19 availability on tumor cells’ surfaces.

We recently established VL51-Ibru, an ibrutinib-resistant model derived from the splenic marginal zone lymphoma (MZL) cell line VL51 through prolonged drug exposure ^8^. Analysis of RNA-sequencing data revealed a significant upregulation of CD19 transcripts in VL51-Ibru compared with parental cells (Figure 1A). Flow cytometry confirmed a corresponding increase in surface CD19 protein expression (Figure 1B). We hypothesized that prolonged exposure to ibrutinib had induced CD19 upregulation due to epigenetic remodelling. To test this hypothesis, we first analysed the DNA methylation status of the CD19 promoter in VL51 exposed or not to ibrutinib. The analysis revealed that the CD19 promoter was hypomethylated in VL51-Ibru cells (beta 0.186) while hypermethylated in parental VL51 cells (beta 0.887, p=0.006; Figure 1C). We next analyzed chromatin immunoprecipitation sequencing (ChIP-seq) data obtained in VL51-Ibru and parental VL51 cells ^9^, focusing on trimethylation of histone H3 at lysine 4 (H3K4me3) associated with active promoters. VL51-Ibru cells exhibited increased H3K4me3 enrichment at the CD19 promoter compared with parental VL51 (Figure 1D). Consistently, assay for transposase-accessible chromatin sequencing (ATAC-seq)^8^ demonstrated enhanced chromatin accessibility at both the CD19 promoter and distal enhancer regions in VL51-Ibru cells (Figure 1E–F). These data indicate that prolonged ibrutinib exposure induced epigenetic remodelling of the CD19 locus, resulting in sustained transcriptional and surface upregulation of CD19 in VL51 cells. To validate our findings in additional B-NHL models, we screened a panel of diffuse large B-cell lymphoma (DLBCL) cell lines and identified two models, DB and SU-DHL-2, that exhibited low baseline CD19 expression. We evaluated the effect of ibrutinib exposure at various concentrations on CD19 surface levels. One-week exposure to ibrutinib (5μM) upregulated CD19 expression on both DB and SU-DHL-2 cells (Supplemental Figure 1A–B). CD19 modulation was dose-dependent, with significant upregulation observed starting at 100 nM in DB cells and 1 μM in SU-DHL-2 cells (Supplemental Figure 1C-D). In contrast, other BTK inhibitors, including acalabrutinib, zanubrutinib, and pirtobrutinib, induced only a minor upregulation of CD19 in DB but not SUDHL-2 cells (Supplemental Figure 2).

**Figure 1.**
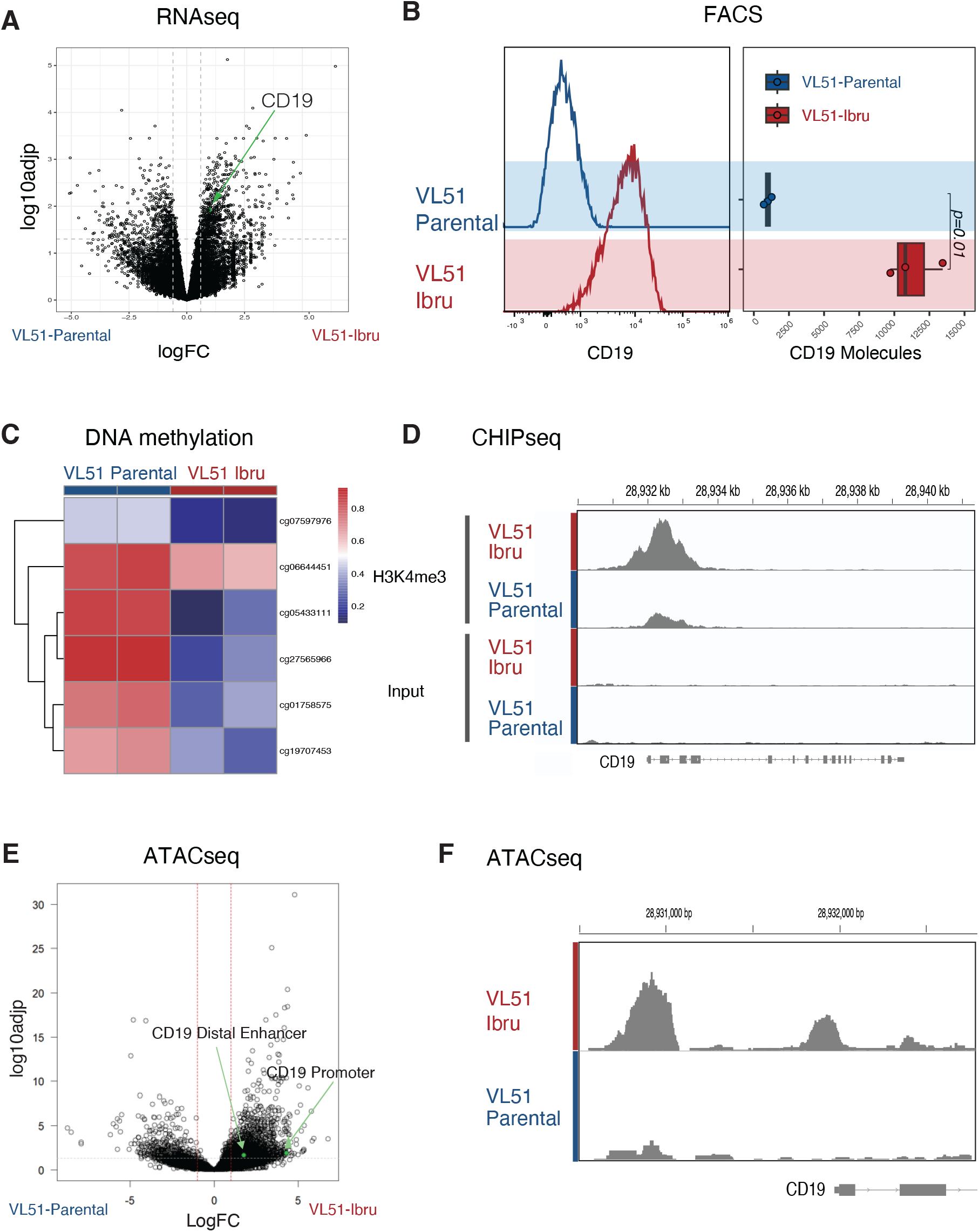
Epigenetic remodeling drives CD19 upregulation in ibrutinib-treated VL51 cells. (A) Volcano plot showing significance and log_2_ fold change of RNA transcripts differentially expressed between VL51-Ibru and parental VL51 cells. (B) Flow cytometry plot of CD19 expression (left panel) and CD19 surface molecule quantification (right panel) of parental VL51 (blue) and VL51-Ibru (red) cells. (C) Heatmap representing DNA methylation levels (β -values) at six specific CpG sites within the CD19 promoter region across two biological replicates. The color scale ranges from blue (hypomethylation, β ≤ 0.2) to red (hypermethylation, β ≥ 0.8). (D) Genome track of H3K4me3 level by ChIP-seq in both ibrutinib-resistant and parental VL51. (E) Volcano plot showing significance and log_2_ fold change of ATAC-seq peaks between VL51-Ibru and parental VL51. (F) Genome tracks showing a comparison of chromatin accessibility by ATAC-seq for the CD19 promoter of VL51-Ibru and parental VL51.

We next investigated the functional consequences of CD19 upregulation on CAR T-cell activity. Human CD19-directed CAR T cells were generated from healthy donor peripheral blood mononuclear cells using retroviral transduction (Online Supplemental Methods). In co-culture assays, longitudinal tumor cell growth was monitored using live-cell imaging (IncuCyte) (Online Supplemental Methods). CD19-CAR T cells exhibited limited cytotoxicity against parental VL51 cells, consistent with their low baseline CD19 expression, while VL51-Ibru cells displayed markedly increased sensitivity to CAR T-cell–mediated killing (Figure 2A-B). Similarly, VL51-Ibru cells exhibited increased, dose-dependent sensitivity to the CD19-targeted antibody-drug conjugate loncastuximab tesirine compared with parental VL51 (Supplemental Figure 3), further supporting the functional relevance of CD19 upregulation across CD19-directed therapies. We next assessed the ability of parental and VL51-Ibru cells to activate CAR T cells. Consistent with enhanced target recognition, CAR T cells co-cultured with ibrutinib-exposed tumor cells produced significantly higher levels of effector cytokines, including IL-2, IFN-γ, and TNF-α (Figure 2C), as measured by multiplex bead-based assays (Online Supplemental Methods).

**Figure 2.**
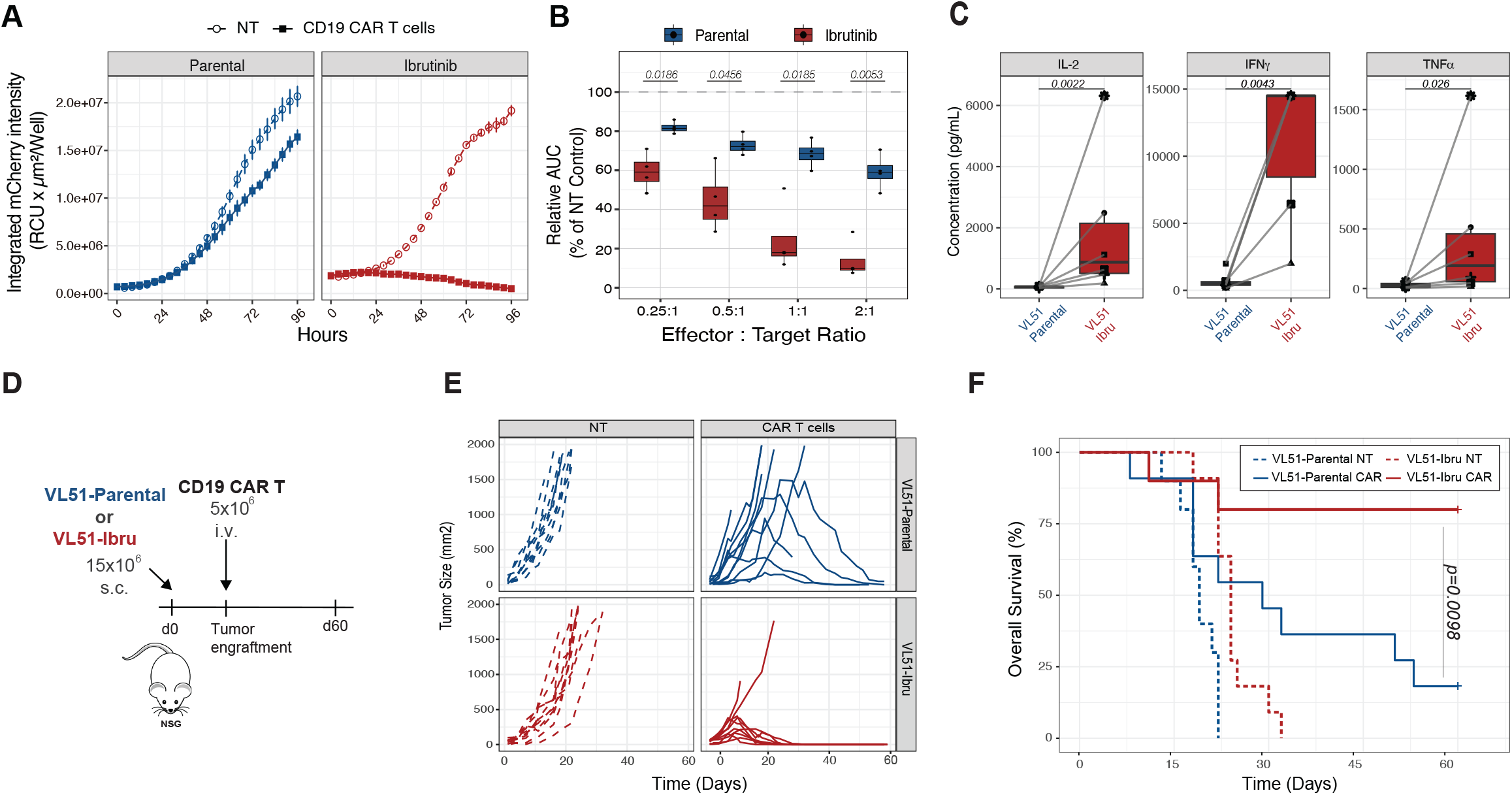
CD19 upregulation led to increased sensitivity to CD19 CAR T cells *in vitro* and *in vivo*. (A) Longitudinal tumor growth of parental VL51 (blue) and VL51-Ibru (red) cells in the presence or absence of CD19 CAR T cells at 1:1 E:T ratios. (B) Area Under the Curve (AUC) analysis normalized by NT tumor cells of CD19 CAR T anti-tumor effect in parental (blue) and ibrutinib (red)-treated VL51 cells. Data are pooled from CD19 CAR T donors (n=4). AUC: area under curve. (C) Analysis of cytokines (IL-2, IFN-g and TNF-a) secreted by CAR T cells derived from 6 donors after 48 hours of co-incubation with either parental VL51 or VL51-Ibru cells at E:T ratio 1:1. (D) Schematic representation of the in vivo experiment. (E) Tumor size from non-treated (NT) and CD19 CAR T groups in both parental (upper panel) and ibrutinib-resistant (lower panel) VL51 tumor bearing mice. (F) Survival curves of mice receiving parental VL51 (blue lines) and VL51-Ibru (red lines) cells treated with CD19 CAR T cells (solid lines) or left non-treated (NT) (dash lines). Results are pooled from two independent experiments with a total of 10 to 11 mice per group. Survival curves were plotted using the Kaplan–Meier method and compared by log-rank test. P values are indicated when significant.

The impact of CD19 modulation was further validated *in vivo*. Immunodeficient NSG mice were subcutaneously engrafted with parental or VL51-Ibru cells and treated with intravenous CD19-CAR T cells following tumor establishment (Figure 2D; Online Supplemental Methods). Tumor growth and survival were monitored longitudinally. CAR T cells induced only partial and delayed tumor control in mice engrafted with parental VL51 cells (Figure 2E). In contrast, CAR T-cell treatment resulted in complete tumor eradication in most mice bearing VL51-Ibru cells (Figure 2E), translating into significantly prolonged survival (Figure 2F). These findings demonstrate that pharmacologic upregulation of CD19 can convert a poorly responsive tumor into one that is highly susceptible to CAR T-cell therapy.

Collectively, our data identify antigen density as a pharmacologically tractable determinant of CAR T-cell efficacy in a B-NHL model. Ibrutinib induced epigenetic remodeling of the CD19 locus, leading to increased antigen expression and enhanced CAR T-cell activation and cytotoxicity.

Several preclinical and clinical studies have reported that ibrutinib may enhance CAR T-cell therapy ^10^ through multiple mechanisms, including improvement of T-cell fitness and product quality ^11–13^, enhanced antitumor activity ^14,15^, reduction of T-cell exhaustion ^16^, modulation of the tumor microenvironment ^16^, and attenuation of treatment-related toxicities ^17^. Our findings identify an additional, previously underappreciated mechanism by which ibrutinib may optimize CD19 CAR T-cell efficacy, namely the modulation of target antigen density at the tumor cell surface. Combination of ibrutinib with CD19 CAR T cells has been tested in clinical trials and appears feasible ^18,19^, although a significant fraction of patients required discontinuation due to toxicities. Our data support the concept of pharmacological antigen priming before CAR T-cell therapy to optimize their efficacy. Short-term exposure to ibrutinib prior to CAR T-cell infusion could enhance tumor susceptibility while avoiding the toxicities associated with concurrent administration.

The use of more selective BTK inhibitors could mitigate toxicities associated with the broader kinase inhibition profile of ibrutinib. However, consistent with prior reports of ibrutinib–CAR T synergy ^12,20^, second-generation BTK inhibitors showed substantially weaker effects on CD19 upregulation, suggesting that this phenotype depends on inhibition of kinases other than BTK, uniquely targeted by ibrutinib. Defining the kinases and signaling pathways regulating CD19 expression may identify safer and more effective therapeutic strategies.

In conclusion, we demonstrate that ibrutinib enhances CAR T-cell efficacy by increasing CD19 expression through epigenetic mechanisms. Pharmacologic modulation of antigen density represents a promising strategy to overcome tumor-intrinsic resistance and improve outcomes following CAR T-cell therapy.

## Supplemental Figure Legends

**Supplemental Figure 1. Ibrutinib induced CD19 upregulation in other B cell lymphoma cell lines**. (A-B) CD19 staining of parental (blue) and 5μM ibrutinib-treated (red) DB and SUDHL2 cells and isotype control (black). (C-D) CD19 surface molecule quantification of DB and SUDHL2 in the presence of different of Ibrutinib after 7 days.

**Supplemental Figure 2. Impact of acalabrutinib, zanubrutinib, and pirtobrutinib on CD19 expression on DB and SUDHL-2 cells**. CD19 surface molecule quantification of DB (upper panel) and SUDHL2 (lower panel) at different concentrations of acalabrutinib (left column), zanubrutinib (middle column) and pirtobrutinib (right column).

**Supplemental Figure 3. Ibrutinib-resistant VL51 cells displayed increased sensitivity to anti-CD19 antibody-drug conjugate loncastuximab tesirine (ADCT-402)**. Viability of parental VL51 (blue) and VL51-Ibru (red) cells in the presence of different doses of ADCT-402.

