## Supplemental Methods for "Ibrutinib modulates CD19 expression and improves efficacy of CD19-CAR T cells in B-cell lymphoma models"

### Online Supplemental Methods

#### Cell lines

B-NHL cell lines (VL51, DB, SU-DHL-2) were cultured in RPMI 160 media supplemented with fetal bovine serum (FBS) and 1% penicillin-streptomycin. Ibrutinib-resistant VL51 cells (VL51-Ibru) were previously reported [1].VL51 parental cell line was transduced with a retroviral vector encoding CD19 (MSCV-hCD19 Plasmid #127889, Addgene VL51 parental, VL51 ibrutinib-resistant and VL51-CD19OE cells were stably transduced with mCherry/luciferase (Plasmid #75020, Addgene).

#### Drug treatments

DB and SU-DHL-2 cells were seeded in culture with DMSO or different concentrations of BTK inhibitors at various concentrations (10 µM, 5 µM, 1 µM, 0.5 µM, and 0.1 µM). The cells were resuspended in fresh medium containing the drugs at day 3 or 4 and analyzed by flow cytometry at day 7. Ibrutinib was obtained from Selleckchem, acalabrutinib, zanubrutinib, and pirtobrutinib from MedChemExpress, and Loncastuximab tesirine was kindly provided by ADC Therapeutics.

#### CD19 expression quantification by flow cytometry

CD19 expression was assessed using PE/APC anti-CD19 antibody (392506/302212, Biolegend) alongside their isotype control (400112/400122, Biolegend). Dead cells were excluded from analysis using DRAQ7 dye (ThermoFisher). The surface expression density of CD19 on various cell lines was determined through saturation binding with the BD Quantibrite Beads PE fluorescence quantitation kit (340495, BD Biosciences) and PE-conjugated anti-CD19 antibody (392596, Biolegend), following the manufacturer's instructions. Maximum antibody binding values and calibration beads were used to quantify the number of CD19 antibody binding sites per cell. For density antigen calculation, PE-labeled calibration beads represent four populations with different numbers of molecules. Using known PE to antibody ratios, the number of PE molecules per cell was converted into the number of antibodies bound per cell. FlowJo software version 10.10.0 was employed for flow cytometry analysis.

#### DNA Methylation

Methylation profiling was performed using the MethylationEPIC BeadChip Infinium, following the manufacturer's instructions for automated array processing with a liquid handler

(Illumina Infinium HD Methylation Assay Experienced User Card, Automated Protocol 15019521 v01), as previously described [2].

**CHIP-seq**

ChIP-seq was performed using a H3K4me3 antibody (17-711 614, RRID:AB_11212770, Millipore) or control IgG (12-371, RRID:AB_145840, Millipore) as negative control. In brief, 50 million cells were resuspended and fixed with 1% methanol-free formaldehyde. Nuclei were extracted, washed and resuspended 25 million/ml in sonication buffer (10 mM Tris pH 8.0, 100 mM NaCl, 1 mM EDTA, 0.5 mM EGTA, 0.5% N-lauroylsarcosine sodium salt and fresh 0.1% Na715 deoxycholate and proteinase inhibitors). Chromatin was fragmented by sonication with Covaris to an average size of 200-500 bp. A 50 µL aliquot was saved as input, whereas 650 µL (corresponding to 16 millions of nuclei) each were incubated with 10 µg antibody, overnight at 4°C. DNA-protein complex was recovered overnight at 65°C, then proteins and RNA were digested and DNA purified by PCR purification kit (Qiagen). Library preparation from 5 ng purified chromatin was performed using NebNext Ultra II DNA Library Prep with magnetic purification beads (E7103S, New England Biolabs) and multiplex oligonucleotides (Dual Index Primers; E7600S, New England Biolabs), followed by adaptor ligation according to the manufacturer's protocol. Quality controls of amplified chromatin fragments were performed on Bioanalyzer 2100 (Agilent Technologies) and Qubit V4 (Thermo Fisher Scientific). Next-generation sequencing was performed on a NextSeq 2000 (Illumina) using the P2 reagent kit V3 (100 cycles; 726 Illumina). Samples were processed starting from stranded, single-ended 120bp-long sequencing reads. Raw data in Fastq format were first preprocessed to ensure suitability for downstream analysis. Adapter sequences were trimmed from the reads using trimmomatic algorithm. Quality control was performed to filter out reads with low-quality bases, short lengths, or those suspected to be sequencing artifacts. Reads were mapped to the HG38 human genome using BWA alignment. Peak calling was performed using MACS (version 2) with "broadPeak" setting, then annotated to the nearest genes with HOMER considering both transcription start and end sites. Raw and processed data have the accession number (pending) and were uploaded on Gene Expression Omnibus (link pending).

**Assay for Transposase-Accessible Chromatin with sequencing (ATAC-seq)**

ATAC-seq for cell lines was performed following the Omni-ATAC-seq protocol 24 with some modifications. Fifty thousand viable cells were lysed in 10 mM tris-HCl (pH 7.4), 10 mM NaCl, 3 mM $MgCl_2$, and 0.1% Igepal CA-630 (Nonidet P-40). The nuclear pellet was then subjected to transposition reaction in 50 μL volume using Illumina Tagment DNA Enzyme and Buffer Small Kit (catalogue n. 20034197) at 37°C for 30 minutes. Then 9 μL per well of stopping mix was added (Clean up buffer 5 μL (900 mM NaCl and 30 mM EDTA), 5% SDS 2 μL and Proteinase K (20 mg/ml) 2 μL) and then incubated at 40°C for 30 min. DNA was then cleaned up with Agencourt AMPure XPBeckman Coulter. PCR amplification was performed according to the protocol for OneTaq Hot Start 2X Master Mix with GC Buffer (M0485, New England Biolabs). The libraries were cleaned up with Agencourt AMPure XP- Beckman Coulter.

**Human CAR T cell generation**

Human peripheral blood mononuclear cells (PBMCs) were collected from the buffy coat of healthy donors (provided by the University Hospitals of Geneva (HUG), Switzerland) using Ficoll-Paque Plus (Cytiva). The cells were then frozen in aliquots and stored in liquid nitrogen. Cryopreserved PBMCs were thawed and activated for 3 days with Dynabeads human T-Activator CD3/CD28 (11132D, Gibco) in the presence of recombinant human IL-2 (50 U/mL, PeproTech) in RPMI1640 media supplemented with 10% FBS and 1% penicillin-streptomycin. Activated T cells were then transduced by culturing them for 48 hours in retronectin (T100B, Takara)-coated 24-well plates with retrovirus encoding second-generation anti-CD19-CD28-CD3ζ CAR. Dynabeads were then removed, and cells were cultured in complete media with 100 U/mL IL-2. Transduction efficacy of CAR expression on T cells was evaluated by APC/PE-conjugated streptavidin (405207/405204, Biolegend) binding to biotinylated protein L (29997, ThermoFisher) or CD19 protein (130-129-550, Miltenyi). At day 14, cells were harvested and frozen for use.

***In vitro* killing assay**

Cell lines transduced with mCherry-luciferase reporter gene were cocultured with either CD19 CAR T cells or untransduced T cell counterparts at different effector/target ratios (E:T) starting from 2:1. CAR T cell number was determined based on transduction efficacy. Non-treated tumor cells were also seeded as a control. The real-time cell growth of all cell lines was measured using mCherry and the Incucyte S3 live-cell analysis instrument (Sartorius).

***In vitro* cytokine detection assay**

Cell lines were cocultured with CD19 CAR T cells at E:T of 1:1. At 48 hours, supernatants were collected and analyzed for cytokine detection (IL-2, IL-6, IL-10, IFN-g, TNF-a) using the LEGENDplex kit (741036, Biolegend) according to the manufacturer's protocols. The

LEGENDplex data analysis software suite was used to analyze LEGENDplex flow cytometry data files.

#### ***In vivo*** **study**

6-8 old weeks NSG mice were subcutaneously engrafted with VL51 parental or ibrutinib-resistant cell line $15x10^6$ cells per mouse. Once the tumor became detectable, CD19 CAR T cells were intravenously injected with $5x10^6$ cells adjusted to transduction. Tumor volume, body weight and general conditions of the mice were monitored twice per week.

#### **Statistical analysis**

Statistical analysis, longitudinal cell growth curves and survival curves were performed using R studio version 2023.12.0.
