## Supplementary figures and images for "Ibrutinib modulates CD19 expression and improves efficacy of CD19-CAR T cells in B-cell lymphoma models"

### Supplemental Figure 1

#### Supplemental Figure 1

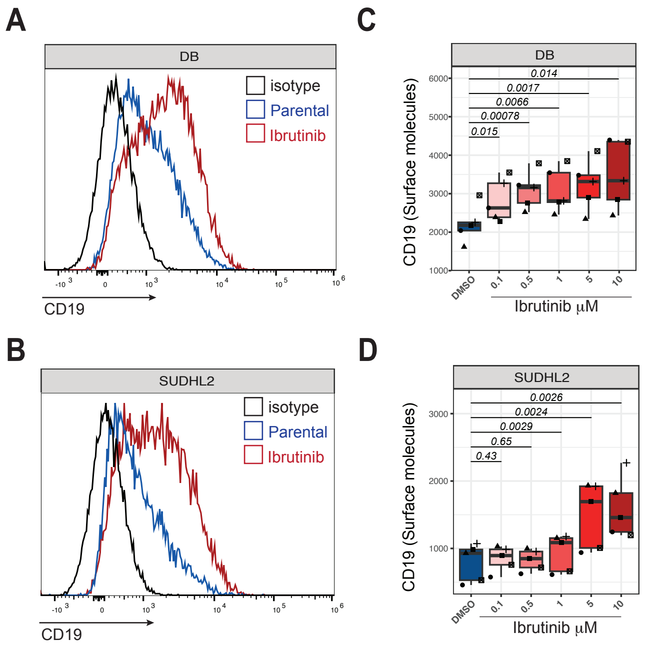

### Supplemental Figure 2

#### Supplemental Figure 2

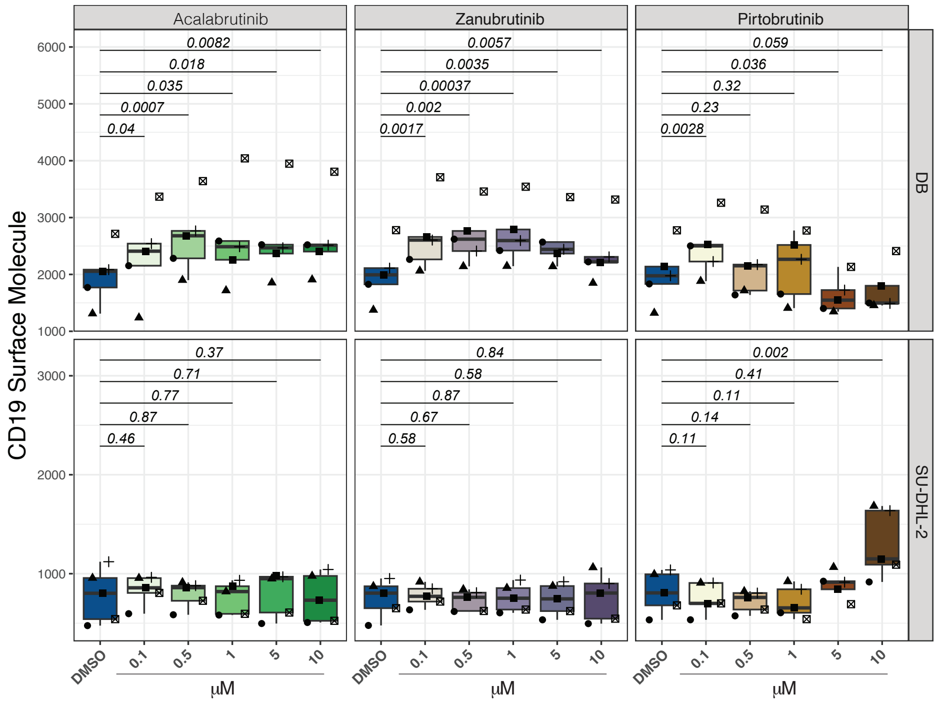

### Supplemental Figure 3

#### Supplemental Figure 3

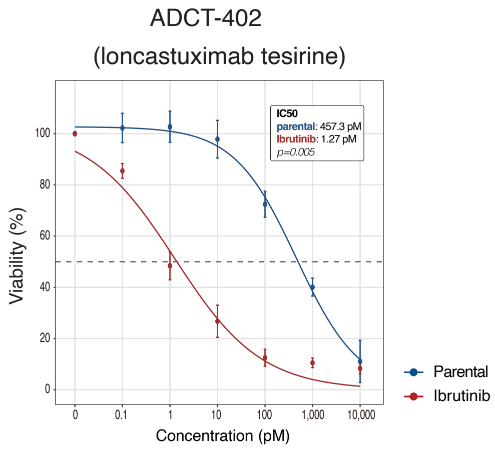
